# The small Canidae from Cuvieri Cave (Quaternary, Pleistocene), Lagoa Santa, eastern Brazil

**DOI:** 10.1101/2025.04.22.650051

**Authors:** Artur Chahud

## Abstract

The Lagoa Santa region is an important karst complex, known for its large number of caves containing abundant fossil and subfossil osteological material. Among the Pleistocene fossils identified in this important area are various remains of paleovertebrates, including members of the order Carnivora. This study describes two fragmented hemimandibles of a small Canidae of the genus *Speothos*, found in Cuvieri Cave. The specimens likely belonged to the same individual, whose second molar had not yet erupted, indicating that it was a subadult. Although the presence of canids has previously been reported in the region, this is the first identification of *Speothos* in the Pleistocene deposits of Cuvieri Cave. It is believed that the animal entered the cave in search of prey, such as rodents and other mammals present at the site, and eventually became trapped within the cavity.

## INTRODUÇTION

The Lagoa Santa region, located in the state of Minas Gerais in eastern Brazil, stands out for its karst complex of great paleontological significance. This environment is characterized by an abundance of osteological materials, which have supported numerous studies over the past years (CHAHUD et al. 2021, FABRIS et al. 2025).

The family Canidae Gray, 1821, belonging to the Order Carnivora, has its earliest fossil records from the Eocene of North America and Europe, and remains present in these regions today. Currently, canids are also found in Africa, Asia, South America, and Australia, the latter likely due to introduction by the first humans who arrived in the region.

In Lagoa Santa, Lund identified, among the Pleistocene fossil remains from the caves of the Rio das Velhas valley, vestiges attributed to various canid species, most of which belong to recent and still extant species. Among the specimens identified, Lund recorded five species of South American canids, including the only known fossils of the genus *Speothos* (LUND, 1839; LUND, 1842; WINGE, 1895; PAULA COUTO, 1979; BERTA, 1984; HANSEN, 2012; RUIZ et al., 2024). Currently, this genus is recognized by two species: one extant, *Speothos venaticus* Lund, 1842, and one extinct, *Speothos pacivorus* Lund, 1839.

The Cuvieri Cave, part of the Lagoa Santa karst complex, has revealed a significant volume of osteological material, providing a basis for various studies on mammal paleontology (MAYER et al., 2016; CHAHUD, 2020; CHAHUD et al., 2023; CHAHUD & OKUMURA, 2021a; CHAHUD & OKUMURA, 2021b; CHAHUD & OKUMURA, 2023). However, until now, no representative of the Canidae family had been conclusively identified in the Pleistocene deposits of the region.

This study describes two small fragmented hemimandibles attributed to a subadult *Speothos*, contributing to the knowledge of this genus in the Pleistocene.

## MATERIALS AND METHODS

Cuvieri Cave is a predominantly horizontal cave comprising three small vertical cavities, designated as *Loci* 1, 2, and 3 (Figure 1), with respective depths of 16 meters, 4 meters, and 8 meters. The materials analyzed in this study originate from *Locus* 3 and have been dated to the Late Pleistocene (MAYER et al., 2016; HADDAD-MARTIN et al., 2017).

**Figure 1.**
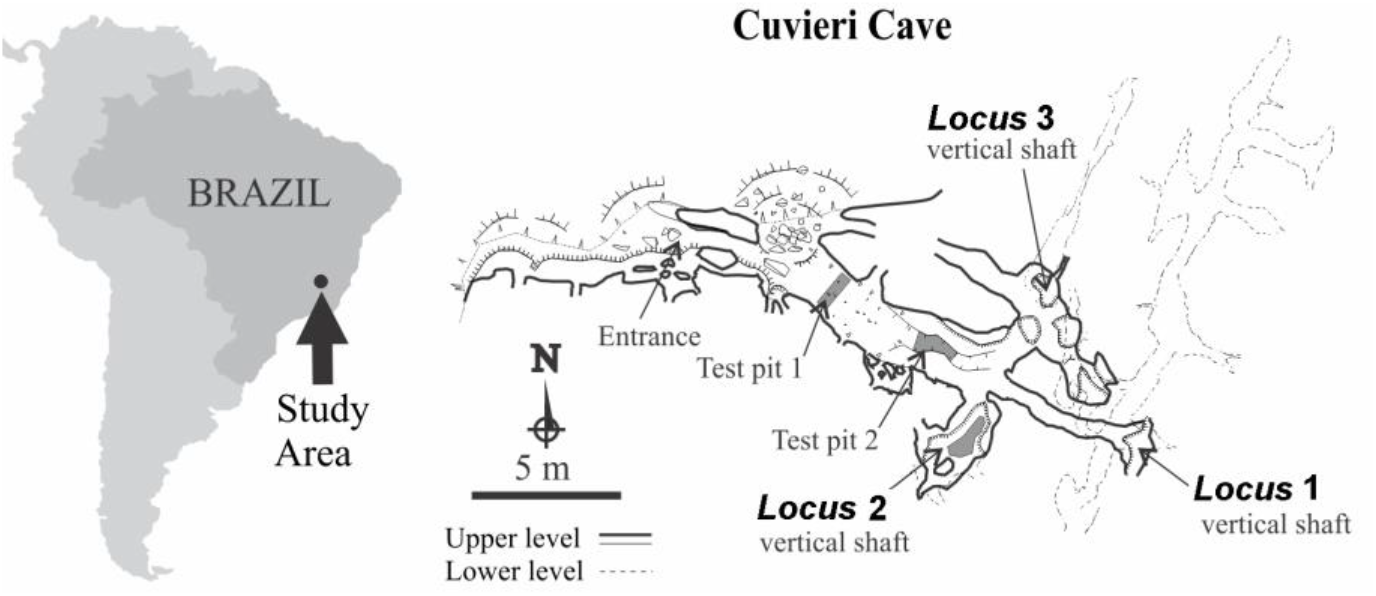
Geographic location of the study area and of Cuvieri Cave showing the position of *Loci* 1, 2 and 3 (map courtesy of Alex Hubbe and Grupo Bambuí for Speleological Research).

The fossil identification was conducted through comparisons with known specimens and consultation of specialized bibliographic references. The two hemimandibles are curated at the Laboratory for Human Evolutionary Studies (LEEH) of the Department of Genetics and Evolutionary Biology at the Institute of Biosciences, University of São Paulo.

### Systematic Paleontology

Order Carnivora Bowdich, 1821

Family Canidae Fischer, 1817

Genus *Speothos* Lund, 1839

*Speothos* sp.

Figures 2 and 3

**Figure 2.**
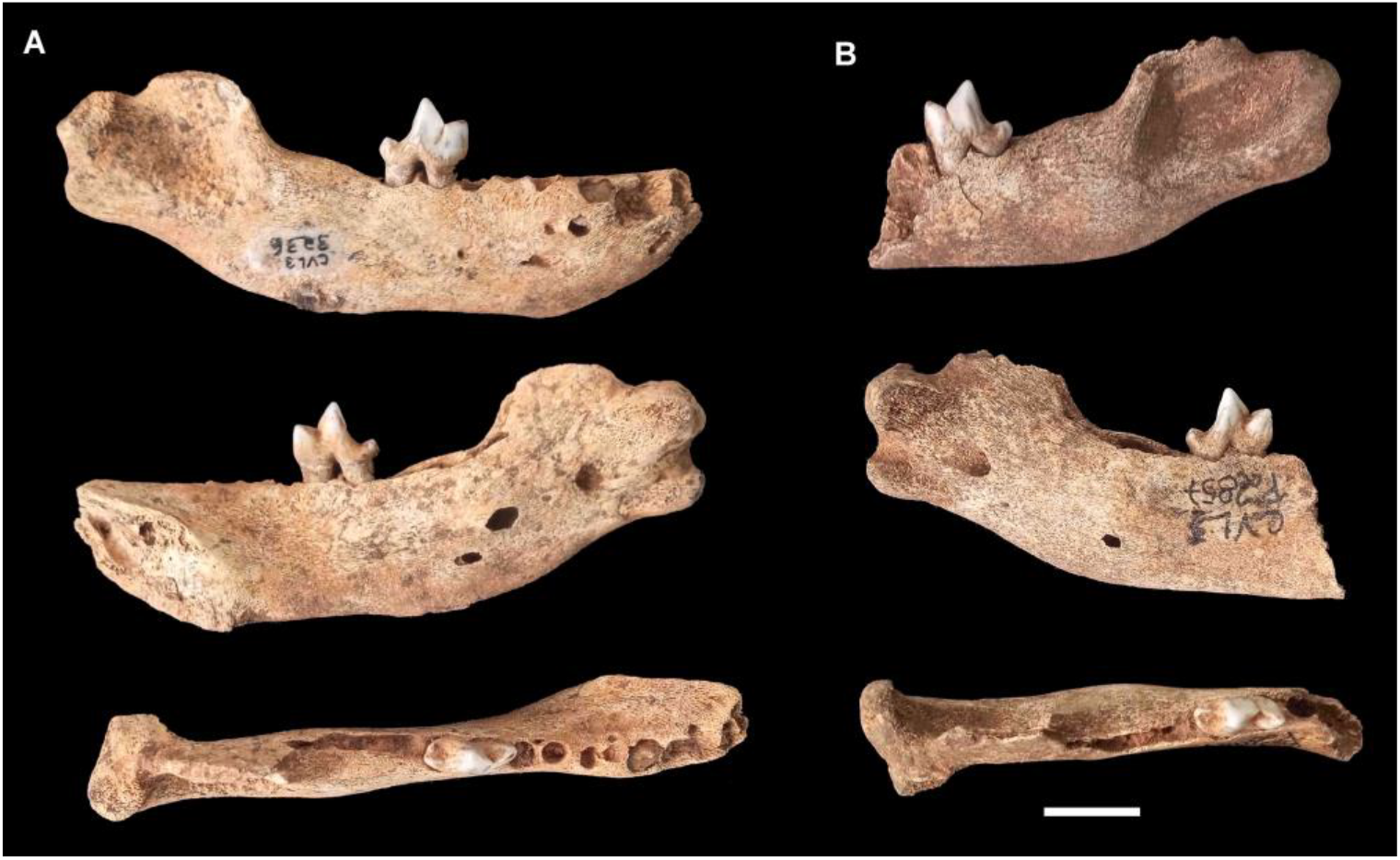
Hemimandibles of *Speothos* from Cuvieri Cave. A) CVL3-3236 (right) and B) CVL3-P2857 (left). From top to bottom: external (labial) view, internal view, and occlusal view. Scale: 10 mm.

**Figure 3.**
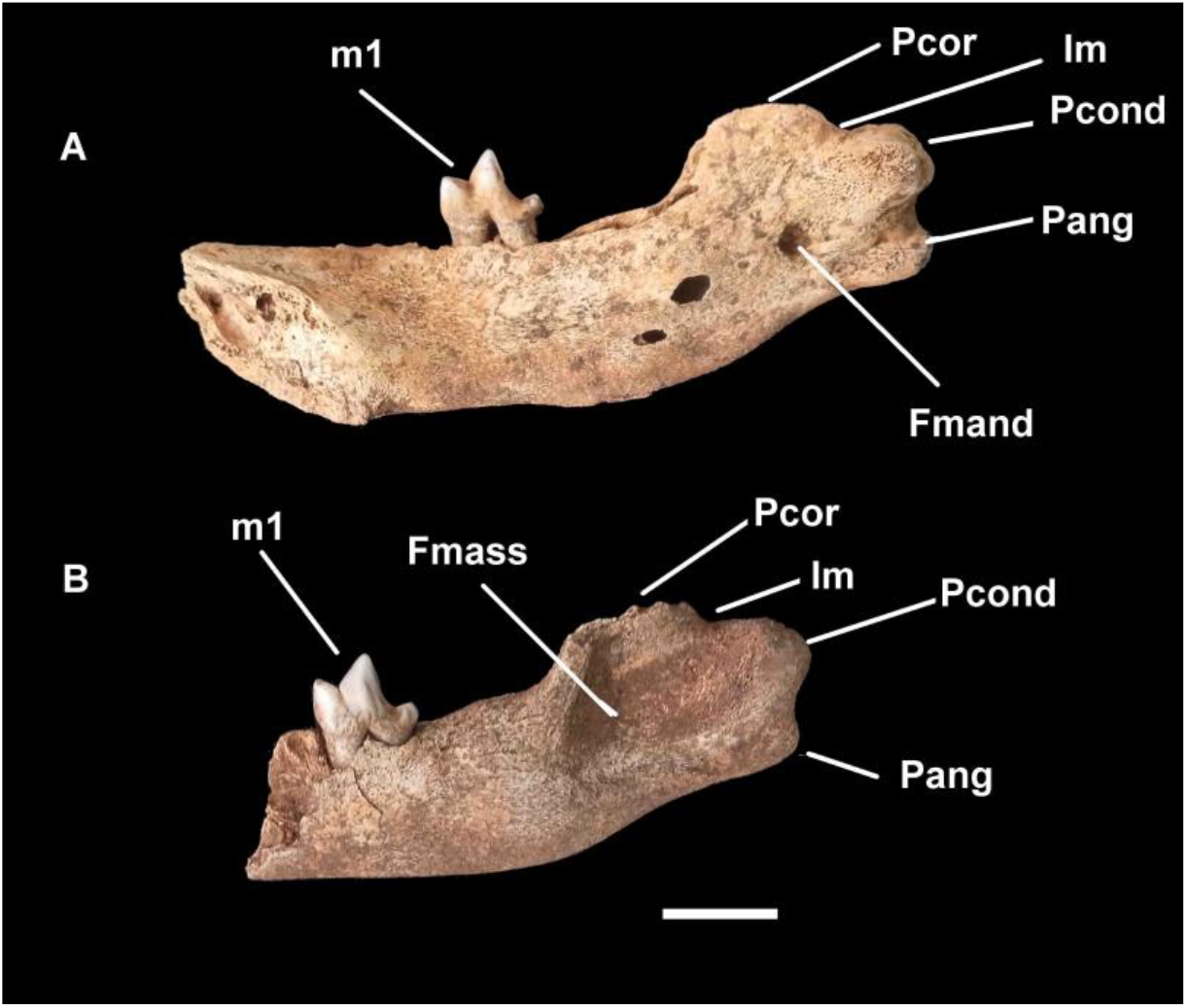
Hemimandibles of *Speothos* from Cuvieri Cave. A) CVL3-3236 (right) and B) CVL3-P2857 (left). **Pcor**: coronoid process; **Im**: mandibular notch; **Pcond**: condylar process; **Pang**: angular process; **Fmand**: mandibular foramen; **Fmass**: masseteric fossa; **m1**: first lower molar. Scale: 10 mm.

### Material

Two fragmented hemimandibles belonging to the same individual, CVL3-3236 (right) and CVL3-P2857 (left).

### Remarks

Both hemimandibles preserve the first molar (m1), part of the mandibular body and ramus, including the masseteric fossa. The morphology of the preserved portions is consistent with that observed in mandibles of the genus *Speothos*, characterized by a short and deep structure compared to other South American canids.

The specimens exhibit prominent mandibular foramina, rounded condylar processes, and relatively small angular processes. The mental foramen is preserved only in hemimandible CVL3-3236 and is located below the alveoli of premolars pm2 and pm3.

The second molar (m2) is absent in both specimens, as is its alveolar cavity, suggesting it had not yet erupted, indicating a subadult individual.

In both specimens, the coronoid process is only partially preserved, having fractured during fossilization while still within the cave. In hemimandible CVL3-3236, the masseteric fossa is worn, and little remains of the condylar and angular processes. In hemimandible CVL3-P2857, the masseteric fossa is more prominent, although the anterior portion of the structure was not preserved.

Molar (m1) measurements: CVL3-3236 (right): width 3.52 mm; length 8.80 mm. CVL3-P2857 (left): width 3.55 mm; length 8.80 mm.

## Discussion

The absence of the second molar (m2) in both specimens suggests that the individual was subadult. However, the masseteric fossa extends close to the first molar (m1), aligning with the expected position of m2, a characteristic also observed in other *Speothos* specimens, both extant and extinct.

The morphology of the mandible, proportionally shorter and deeper than that of other South American canids, supports its attribution to the genus *Speothos*. Additionally, the analyzed hemimandibles are less robust compared to extant specimens, resembling the extinct species *S. pacivorus* (BERTA, 1984; RUIZ et al. 2022; RUIZ et al. 2025). According to RUIZ et al. (2025), modern *Speothos* specimens exhibit proportionally deeper mandibles than *S. pacivorus*, which aligns the Cuvieri Cave specimen more closely with the extinct species. However, the fragmentation of the remains and the subadult condition of the individual may influence this comparison. Therefore, a cautious identification was made, assigning the specimen to *Speothos* sp.

## ASSOCIATED FAUNA

Despite the remobilization of bone material, various mammals were found in the same stratigraphic levels as the analyzed hemimandibles. Among the identified specimens, fossil remains of small rodents, marsupials, Xenarthra, and large rodents such as *Cuniculus paca* (living paca), *C. rugiceps* (extinct paca), and *Dasyprocta* sp. were prominent (MEYER et al. 2016; TAKAHASHI & CHAHUD 2024; 2025), as well as undetermined representatives of Tayassuidae and Tapiridae (CHAHUD et al. 2023).

Large rodents, such as *Cuniculus paca* and *Dasyprocta* sp., are common prey of *Speothos* today. Their frequent occurrence in Cuvieri Cave suggests that the analyzed individual may have been drawn into the cave while pursuing prey. Additionally, remains of extinct megafauna, including ground sloths, were identified.

## FINAL CONSIDERATIONS

The evidence suggests that the two hemimandibles belonged to the same individual, which underwent disarticulation and significant remobilization within Cuvieri Cave. The material corresponds to a juvenile *Speothos* individual, whose second molar had not yet erupted.

Canid representatives have been previously recorded in other studies, although occurrences of *Speothos* are relatively rare. The analyzed individual coexisted with species such as *Cuniculus paca, C. rugiceps, Dasyprocta* sp., Tayassuidae, and Xenarthra and microvertebrates. The exact reason for the animal’s entry into the cave remains uncertain, but it was likely drawn in by potential prey present at the site.

## ACKNOWLEDGEMENTS

The author thanks the Dr. Maria Mercedes Martinez Okumura responsible for LEEH (Laboratory for Human Evolutionary Studies), Department of Genetics and Evolutionary Biology, Institute of Biosciences of the University of São Paulo for permitting the preparation of the fossils in her laboratory.

